# Proteomic Signatures of Mitochondrial Dysfunction Associated with Atrial Fibrillation in Goats

**DOI:** 10.64898/2026.02.23.707419

**Authors:** Thamali Ayagama, Richard Barrett-Jolley, Roman Fischer, Svenja Hester, Ulrich Schotten, Sander Verheule, Rebecca AB Burton

**Affiliations:** Department of Physiology, Anatomy and Genetics, University of Oxford, Oxford, UK; Department of Musculoskeletal & Ageing Science, Institute of Life Course & Medical Sciences Faculty of Health and Life Sciences, University of Liverpool, Liverpool, UK; Target Discovery Institute/Centre for Medicines Discovery, Nuffield Department of Medicine, University of Oxford, Oxford, UK; Departments of Physiology and Cardiology, Cardiovascular Research Institute Maastricht, Maastricht University, Maastricht, The Netherlands; Institute of Systems, Molecular and Integrative Biology, Department of Pharmacology and Therapeutics, University of Liverpool, Liverpool, UK; Liverpool Centre for Cardiovascular Science, University of Liverpool and Liverpool Heart & Chest Hospital, Liverpool, UK; Department of Pharmacology, University of Oxford, Oxford, UK

**Author notes:** Corresponding author / and. Joint first authors (Thamali Ayagama, Richard Barrett-Jolley).

## Abstract

Atrial fibrillation (AF) increases energy demand in atrial myocytes, yet the mitochondrial mechanisms underlying this stress remain poorly defined. Using previously published proteomic data from left atrial tissue of AF and sham-operated goats, we performed organelle-specific bioinformatic analyses of the mitochondrial fraction. Over-representation and consensus pathway analyses consistently highlighted enrichment of oxidative phosphorylation (OXPHOS) subunits. Gene set enrichment and network analyses implicated Heat Shock Protein Family A Member 9 (HSPA9) as a potentially central regulatory hub coordinating the dysregulation of Complex I and III subunits, with 69% of regulatory relationships showing pathway concordance. These results indicate a coordinated, system-wide mitochondrial adaptation in AF, integrating energy production, proteostasis, and respiratory chain regulation.

## Introduction

Atrial fibrillation (AF) is a widespread arrhythmia whose global prevalence has grown substantially in the last three decades, now affecting an estimated 60 million people [1]. Across Europe, prevalent AF cases are projected to grow from about 8.8 million today to nearly 18 million by 2060 [2]. Left unaddressed, AF can cause major complications, including stroke, heart failure, and thromboembolic events, each of which can markedly diminish health and quality of life [3]. A deeper understanding of the molecular mechanisms driving AF is essential for elucidating its pathophysiology and laying the groundwork for precision-medicine strategies in its management [4].

AF is characterized by altered electrical activity and increased energy demands in atrial myocytes [5–7]. Mitochondria, as central players in energy metabolism, generate ATP through oxidative phosphorylation (OXPHOS), a process closely linked to AF pathology [8]. In cardiomyocytes, mitochondrial ATP production supports not only contraction and relaxation but also ion homeostasis, calcium handling, and other energy-intensive processes, making mitochondrial function central to cardiac physiology [9–11]. To meet these demands, the heart relies on an efficient metabolic system in which mitochondria produce ATP through redox reactions coupled to chemiosmotic gradients across their membranes. Beyond ATP generation, mitochondria also regulate cell death and survival pathways and modulate key second messengers, including calcium ions (Ca^2+^) and reactive oxygen species (ROS), further underscoring their multifaceted role in maintaining cardiac function [11].

Mitochondrial dysfunction is a hallmark of numerous cardiovascular diseases, where it contributes to impaired energy metabolism and heightened cellular stress [12, 13]. In heart failure, disturbances in substrate utilization, OXPHOS, and phosphotransfer systems compromise ATP production and energetic homeostasis [14, 15]. Dysregulated mitophagy further accelerates the development of cardiomyopathy and promotes atherosclerosis by enhancing plaque formation and increasing ROS generation from damaged mitochondria [16]. In myocardial infarction and ischemia–reperfusion injury, coronary occlusion induces mitochondrial Ca^2+^ overload and opening of the permeability transition pore, ultimately leading to cell death and extensive tissue damage [17, 18].

Advances in organelle proteomics [19–24] have enabled systematic analysis of organelle specific proteomes such as endolysosomal (EL) proteins [25], allowing identification of differentially expressed proteins and organelle-specific pathway alterations. Coupled with bioinformatic approaches such as network modelling and pathway enrichment analysis, these tools provide mechanistic insights into how organelle dysfunction contributes to cellular stress, energetic remodelling, and AF pathophysiology. This study aims to enhance our understanding of mitochondrial dysfunction in AF by utilizing data obtained from our proteomics study in goat tissue to explore protein regulation within the mitochondrial fraction.

## Methods

### Tissue Homogenisation and Liquid chromatography-tandem mass spectrometry analysis

We previously published proteomic analyses of left atrial tissue from AF and sham goats, in which both tissue lysate (TL) and EL fractions were characterized [26]. The goat study was carried out in accordance with the principles of the Basel declaration and regulations of European directive 2010/63/EU, and the local ethical board for animal experimentation of the Maastricht University approved the protocol. In brief, frozen atrial biopsies (∼100 mg) were washed in phosphate-buffered saline (PBS), minced with sterile scalpels, and homogenized using Dounce homogenizers in lysosome isolation buffer containing protease and phosphatase inhibitors. Homogenates were further processed with lysosome enrichment buffer, followed by sequential centrifugation steps to isolate the TL, EL and mitochondrial fractions. TL was collected after centrifugation at 13,000 × g for 2 min at 4°C. The EL fraction was obtained using a sucrose-Percoll density gradient, with ultracentrifugation at 67,000 × g for 30 min at 4°C, isolating the fraction at 1.04 g/mL. The mitochondrial fraction was obtained by further processing the collected TL at the end of 13,000 g centrifugation. For mitochondrial isolation, 1.5 mL Beckmann ultracentrifuge tubes were underlaid with 750 μL of 2.5 M Sucrose (Fisher Scientific), followed by layering 250 μL of Percoll (Santa Cruz Biotechnology) on top. Subsequently, 200 μL of TL was carefully layered on top of the Percoll layer and centrifuged at 27,000 g × 50 min at 10°C. After centrifugation, a turbid white line became visible at the interface between Percoll and 2.5 M sucrose (at 1.3g/mL), representing an accumulation of mitochondria with some sarcoplasmic reticulum (SR). This fraction was collected into a low protein-binding Eppendorf tube. Three biological replicates per condition were used (n=3). Liquid chromatography-tandem mass spectrometry analysis was performed at the Target Discovery Institute, Oxford. Raw mass spectrometry data were processed in Progenesis QI (WatersTM Cooperation, www.nonlinear.com, v4.2) and searched against the UniProt *C. hircus* database (UP000291000). Human UniProt gene symbols associated with each goat protein (via GN fields) were used for downstream pathway enrichment. Differential abundance was calculated by the Progenesis software using Student’s t-test. Mitochondrial data were submitted to **PRIDE (PRIDE ID: PXD041056)**. In the present study, we leveraged these previously generated proteomics data [26] to perform detailed bioinformatics analyses focusing specifically on the mitochondrial fraction. Supplementary file 1 contains the comprehensive list of quantified mitochondrial proteins identified in the dataset.

### Bioinformatics

#### Proteomics Data Processing and Quality Control

Mitochondrial fraction proteomics data from an AF animal model versus sham controls were analysed using R version 4.5x and Bioconductor packages. Progenesis data (Supplementary file 1) were imported and filtered for unique proteins and entities such as fusion “orf” proteins removed. For proteins identified by multiple peptides, the entry with the highest number of unique peptide hits was retained to ensure confident identification. Very low-abundance proteins (bottom 1st percentile by mean expression) were excluded from downstream analysis.

#### Over-representation Analyses (ORA)

ORA was performed with clusterprofiler [27, 28]. Multiple pathway databases were interrogated to ensure comprehensive functional annotation: Gene Ontology (GO) biological processes (excluding “CC”, cellular compartment), Kyoto Encyclopedia of Genes and Genomes (KEGG) pathways (filtered to remove disease pathways), Reactome pathways [29, 30] and WikiPathways [31]. Default backgrounds were used. Significant protein lists (p ≤ 0.05) were further filtered for absolute log2fold change of greater than 0.65. ORA plots (“dot plots”), category-item networks (CNE), and enrichment maps (EMA) were constructed using enrichplot [32].

#### Cross-Database Consensus Enrichment

To identify robustly enriched pathways independent of database-specific nomenclature or annotation biases, a consensus enrichment approach was implemented across all four of the major pathway databases above. Rather than relying on pathway name matching (which varies substantially across databases), consensus pathways were identified based on gene set overlap. For all pairwise combinations of enriched pathways across databases, Jaccard similarity indices were calculated on the constituent gene sets. Pathways with ≥80% gene overlap (Jaccard index ≥ 0.8) were clustered together regardless of their assigned names in the source database. For each cluster, the quoted Name or Description was selected preferentially from GO, and the number of supporting databases, median *p-value*, and shared gene list were recorded. Database overlap patterns and consensus pathway identification were visualized using an UpSet plot (“pheatmap”)[33].

#### Mitochondrial-specific analyses

Given the mitochondrial-enriched nature of the proteomics fraction, mitochondrial-specific pathways were analyzed using gene set enrichment analysis (GSEA) rather than over-representation analysis to leverage the expected comprehensive coverage of mitochondrial proteins. GSEA was performed using the mitology package and MitoCarta3.0 database [25], a curated compendium of mitochondrial genes representing the gold standard for mitochondrial annotation. Unlike ORA, GSEA utilizes the entire ranked gene list without imposing arbitrary significance thresholds, reducing selection bias and improving statistical power when pathway coverage is expected to be high. We ranked proteins on the bases of LFC weighted *p-values*. Sample-level mitochondrial pathway activities were additionally quantified using single-sample GSEA (ssGSEA) implemented via GSVA [34], and visualized using hierarchical tree heatmaps that organize MitoCarta pathways by their functional relationships

#### Network Analysis

To investigate regulatory relationships within enriched mitochondrial pathways, causal network analysis was performed using the experimentally-validated protein-protein directional interaction data from the Reactome database [35]. Specifically, we imported the “Aerobic respiration and respiratory electron transport” network. For each interaction annotated as “activation” or “inhibition”. Edges were considered concordant when activating proteins and their targets showed co-directional changes. “pathway concordance” *p-value* was calculated by comparing the expected versus observed directional changes in protein expression (with a permutation test).

Two-step pathway chains were identified by joining consecutive edges in the network, and hub proteins (those participating in multiple regulatory cascades) were identified by network degree analysis. Network visualization was performed using the igraph package [36].

## Results

### Overview analysis

Initially, as a positive control, we analysed our mitochondrial fraction data for over-representation (ORA) against the entire genome. Since the data are from a mitochondrial fraction, we would expect this to enrich mitochondrial pathways. We used the Gene Ontology (GO) database for biological processes (BP), molecular function (MF) and cellular compartment (CC) along with KEGG, WikiPathways and Reactome. Genes were defined “DEG” if it’s *p-value* was less than 0.05 and log2 fold-change was greater than 0.65. Significant over-representation was based on Benjamini-Hochberg correction (BH) adjusted *p-values* of 0.05. Figure 1 shows the GO BP dot plot (A) with gene concept network (B) and enrichment map in (C). Figures 2 and 3 show the same information (where calculable) for GO:MF and GO:CC respectively. ORA with the WikiPathways, KEGG and Reactome databases are shown in Figures 4, 5 and 6 respectively.

**Figure 1.**
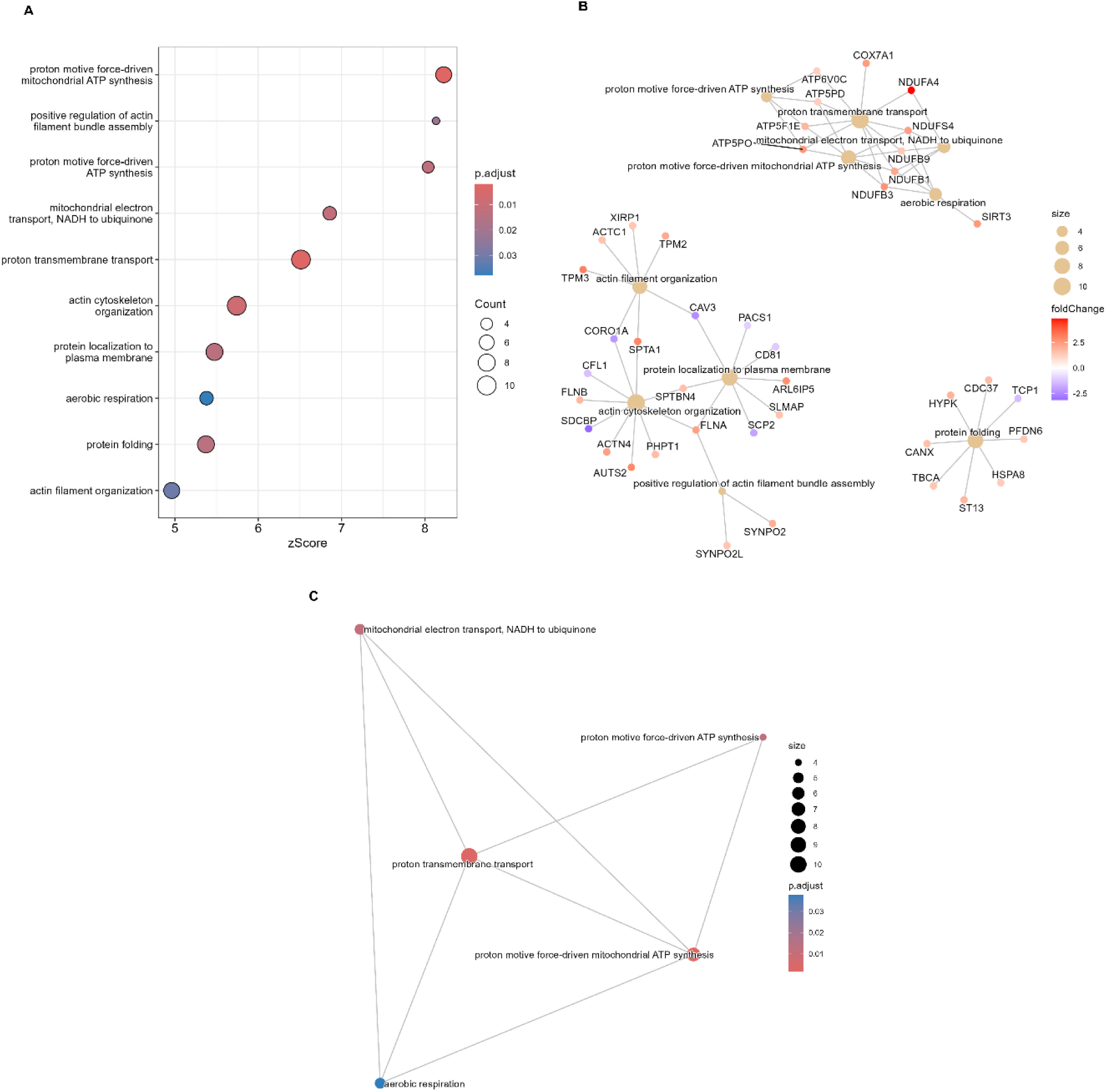
Gene Ontology (GO) Biological Process (BP) enrichment analysis reveals mitochondrial ATP synthesis and respiratory dysfunction in AF samples. (A) Dot plot showing significantly enriched GO BP terms from over-representation analysis of differentially abundant proteins. The x-axis represents enrichment z-score, dot size indicates gene count, and colour intensity represents adj usted p-value (Benjamini-Hochberg correction). Terms are ranked by significance, with proton motive force-driven mitochondrial ATP synthesis showing the highest enrichment. (B) Concept Network Enrichment (CNE) plot displaying gene-concept relationships for top enriched BP. Genes (outer nodes) are connected to their associated GO terms (inner nodes), with node size proportional to gene count or term size. Gene nodes are coloured by log2-fold change (red = upregulated, blue = downregulated in AF). (C) Enrichment Map Analysis (EMA) plot showing functional relationships between enriched biological process terms. Nodes represent GO terms sized by gene count, and edges connect terms sharing genes. Terms cluster by functional similarity, revealing coordinated enrichment of mitochondrial ATP synthesis, electron transport, and actin cytoskeleton organization pathways. All enrichments shown have adjusted p-value < 0.05.

**Figure 2.**
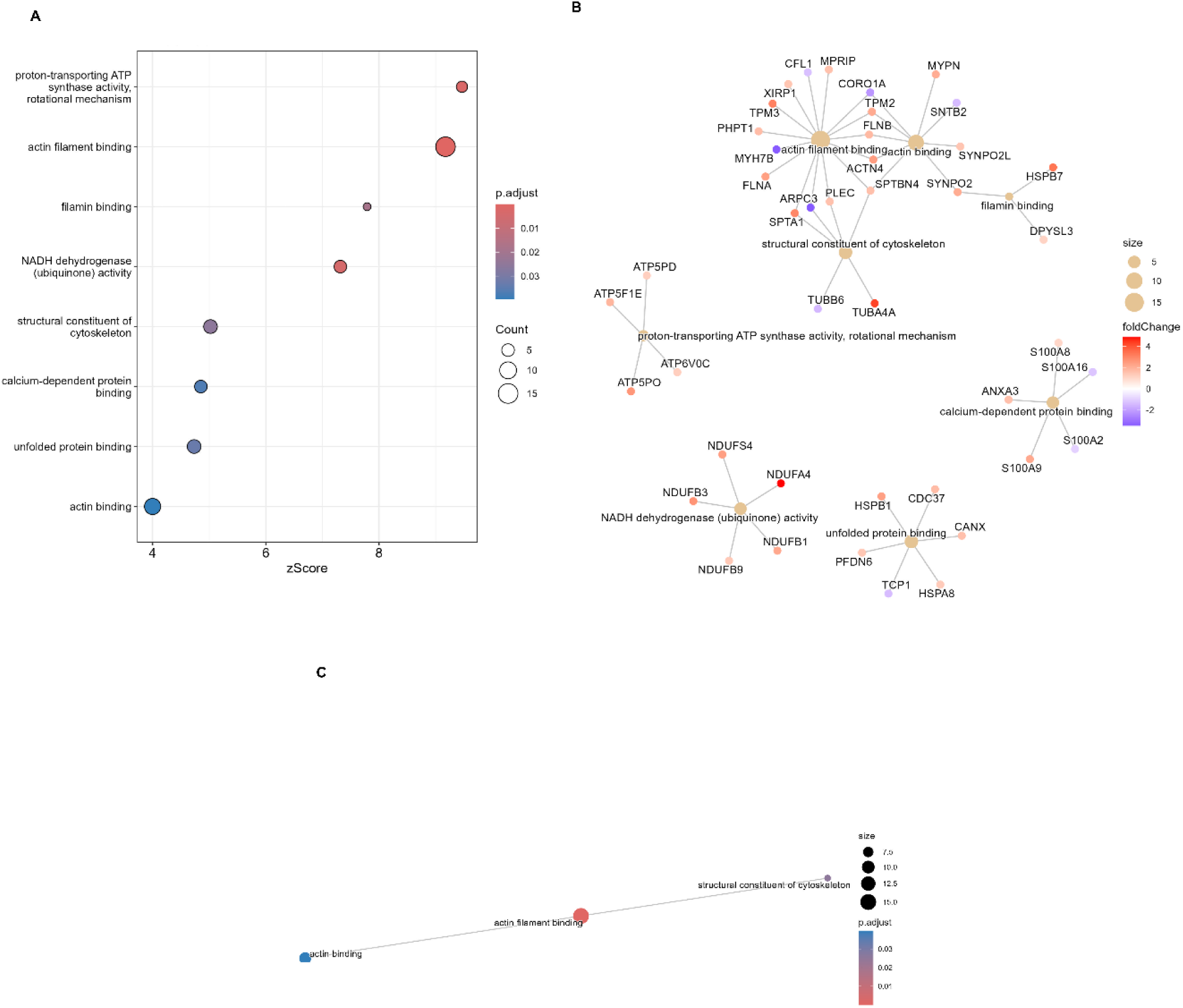
Gene Ontology (GO) Molecular Function enrichment analysis highlights altered ATP synthase and cytoskeletal binding activities in AF samples. (A) Dot-plot showing significantly enriched GO Molecular Function terms from over-representation analysis of differentially abundant mitochondrial proteins. The x-axis represents enrichment z-score, dot size indicates gene count, and colour intensity represents adjusted p-value (Benjamini-Hochberg correction). Proton-transporting ATP synthase activity and actin filament binding show the most significant enrichment. (B) Concept Network Enrichment (CNE) plot displaying gene-concept relationships for top enriched molecular functions. Genes (outer nodes) are connected to their associated GO terms (inner nodes), with node size proportional to gene count or term size. Gene nodes are coloured by log2 fold change (red = upregulated, blue = downregulated in AF). (C) Enrichment Map Analysis (EMA) plot showing functional relationships between enriched molecular function terms. Nodes represent GO terms sized by gene count, and edges connect terms sharing genes. Terms cluster by functional similarity, with prominent groupings of structural constituent of cytoskeleton and actin filament binding functions. All enrichments shown have adjusted p-value < 0.05.

**Figure 3.**
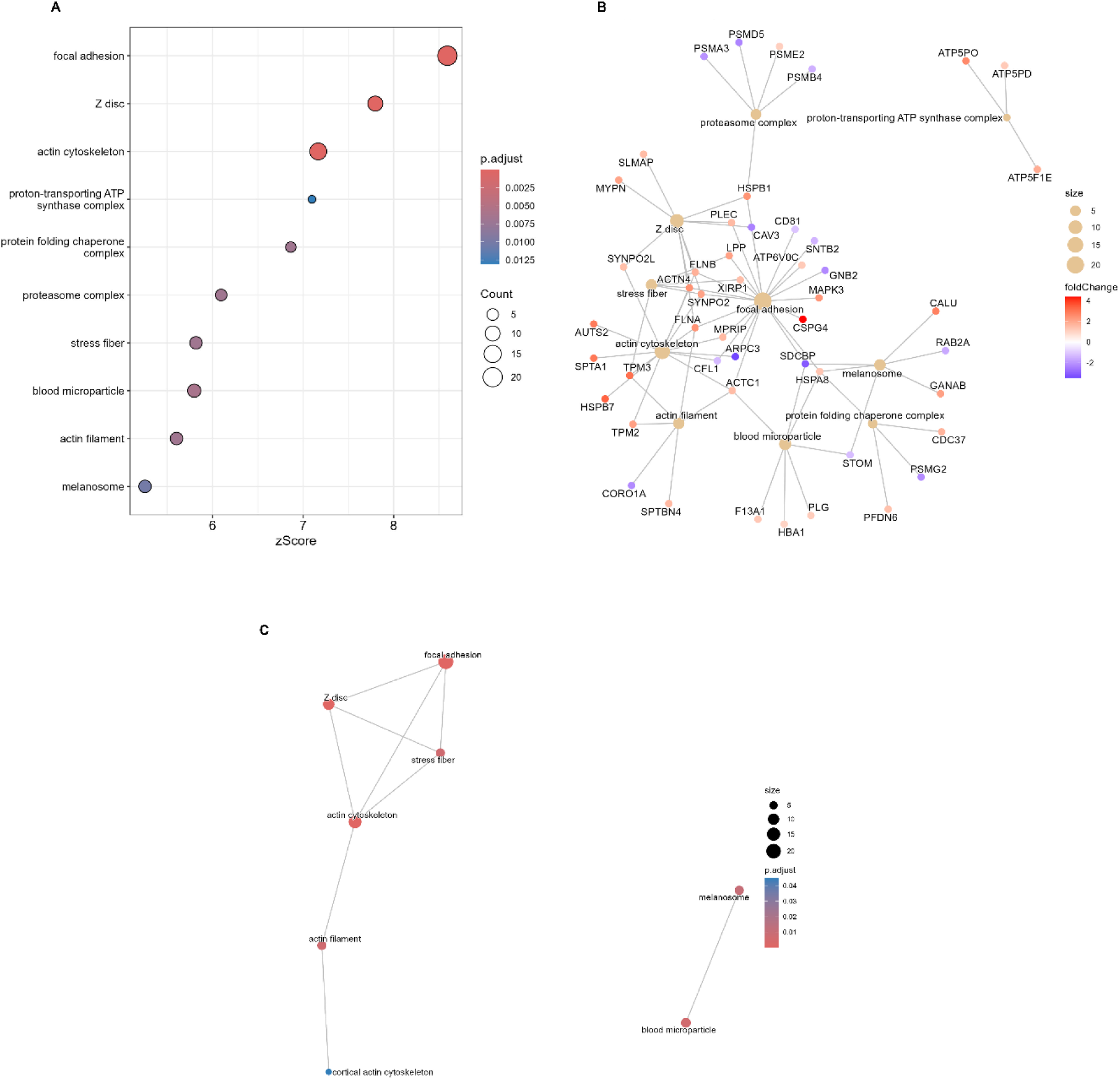
Gene Ontology Cellular Component enrichment analysis reveals reorganization of focal adhesions and mitochondrial complexes in AF samples. (A) Dot-plot showing significantly enriched GO Cellular Component terms from over-representation analysis of differentially abundant mitochondrial proteins. The x-axis represents enrichment z-score, dot size indicates gene count, and colour intensity represents adjusted p-value (Benjamini-Hochberg correction). Focal adhesion, Z disc, and actin cytoskeleton components show the highest enrichment scores. (B) Concept Network Enrichment (CNE) plot displaying gene-concept relationships for top enriched cellular components. Genes (outer nodes) are connected to their associated GO terms (inner nodes), with node size proportional to gene count or term size. Gene nodes are coloured by log2 fold change (red = upregulated, blue = downregulated in AF). (C) Enrichment Map Analysis (EMA) plot showing functional relationships between enriched cellular component terms. Nodes represent GO terms sized by gene count, and edges connect terms sharing genes. Terms cluster into distinct functional modules including focal adhesion/Z disc/actin cytoskeleton, proteasome complex, and blood microparticle compartments. All enrichments shown have adjusted p-value < 0.05.

**Figure 4.**
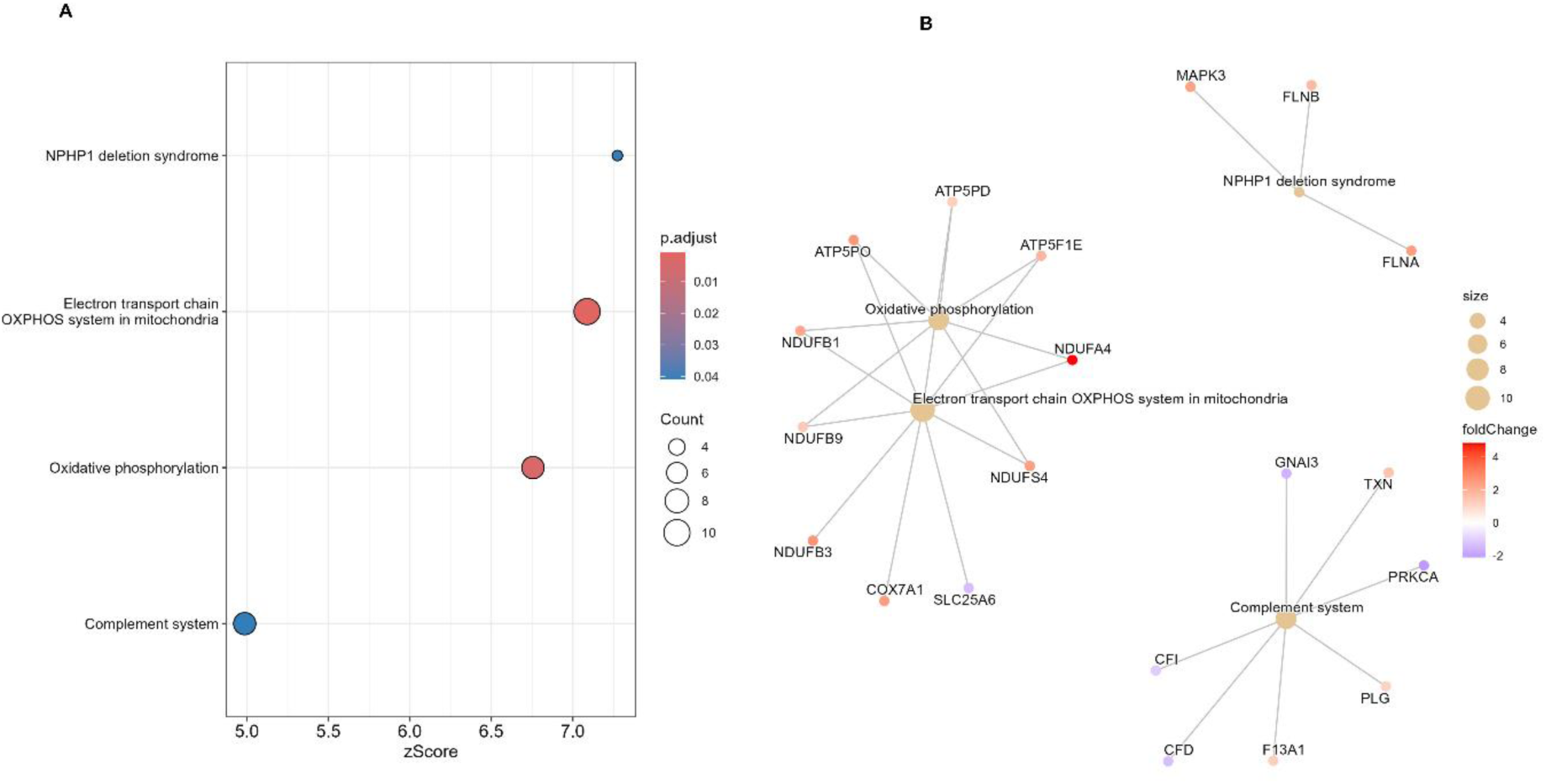
WikiPathways enrichment analysis identifies mitochondrial electron transport chain dysfunction in AF samples. (A) Dot-plot showing significantly enriched WikiPathways terms from over-representation analysis of differentially abundant proteins from AF-induced left atrial tissue versus sham-operated controls. The x-axis represents enrichment z-score, dot size indicates gene count, and colour intensity represents adjusted p-value (Benjamini-Hochberg correction). Electron transport chain OXPHOS system in mitochondria shows the most significant enrichment. (B) Concept Network Enrichment (CNE) plot displaying gene-concept relationships for top enriched pathways. Genes (outer nodes) are connected to their associated pathway terms (inner nodes), with node size proportional to gene count or term size. Gene nodes are coloured by log2 fold change (red = upregulated, blue = downregulated in AF). Electron transport chain and oxidative phosphorylation pathways cluster together with multiple NADH dehydrogenase (NDUF) subunits and ATP synthase components. All enrichments shown have adjusted p-value < 0.05.

**Figure 5.**
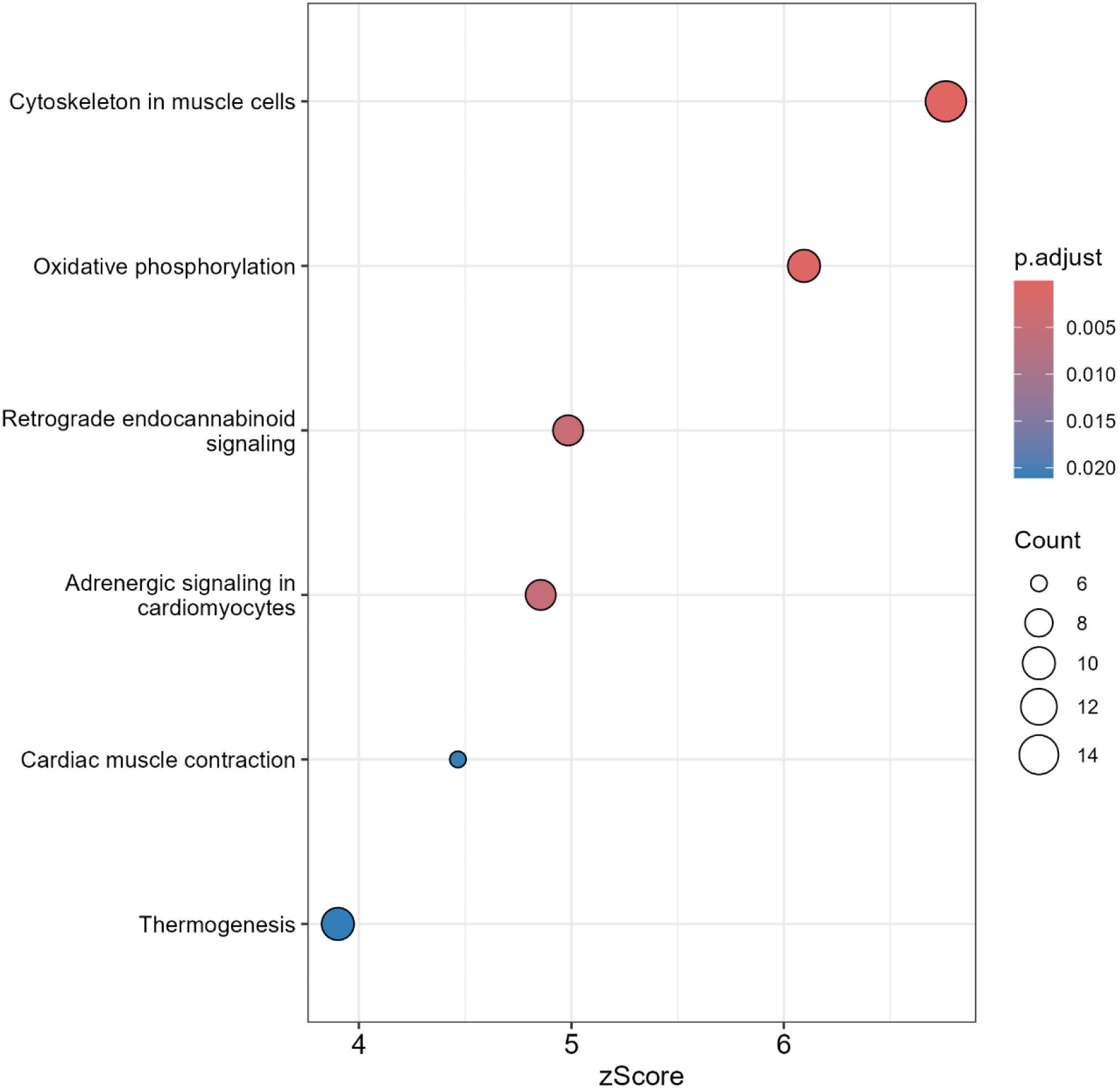
KEGG pathway enrichment analysis reveals muscle-specific and metabolic adaptations in AF samples. (A) Dot-plot showing significantly enriched KEGG pathways from over-representation analysis of differentially abundant proteins from AF-induced left atrial tissue versus sham-operated controls. The x-axis represents enrichment z-score, dot size indicates gene count, and colour intensity represents adjusted p-value (Benjamini-Hochberg correction). Cytoskeleton in muscle cells and oxidative phosphorylation show the highest enrichment scores. All enrichments shown have adjusted p-value < 0.05.

**Figure 6.**
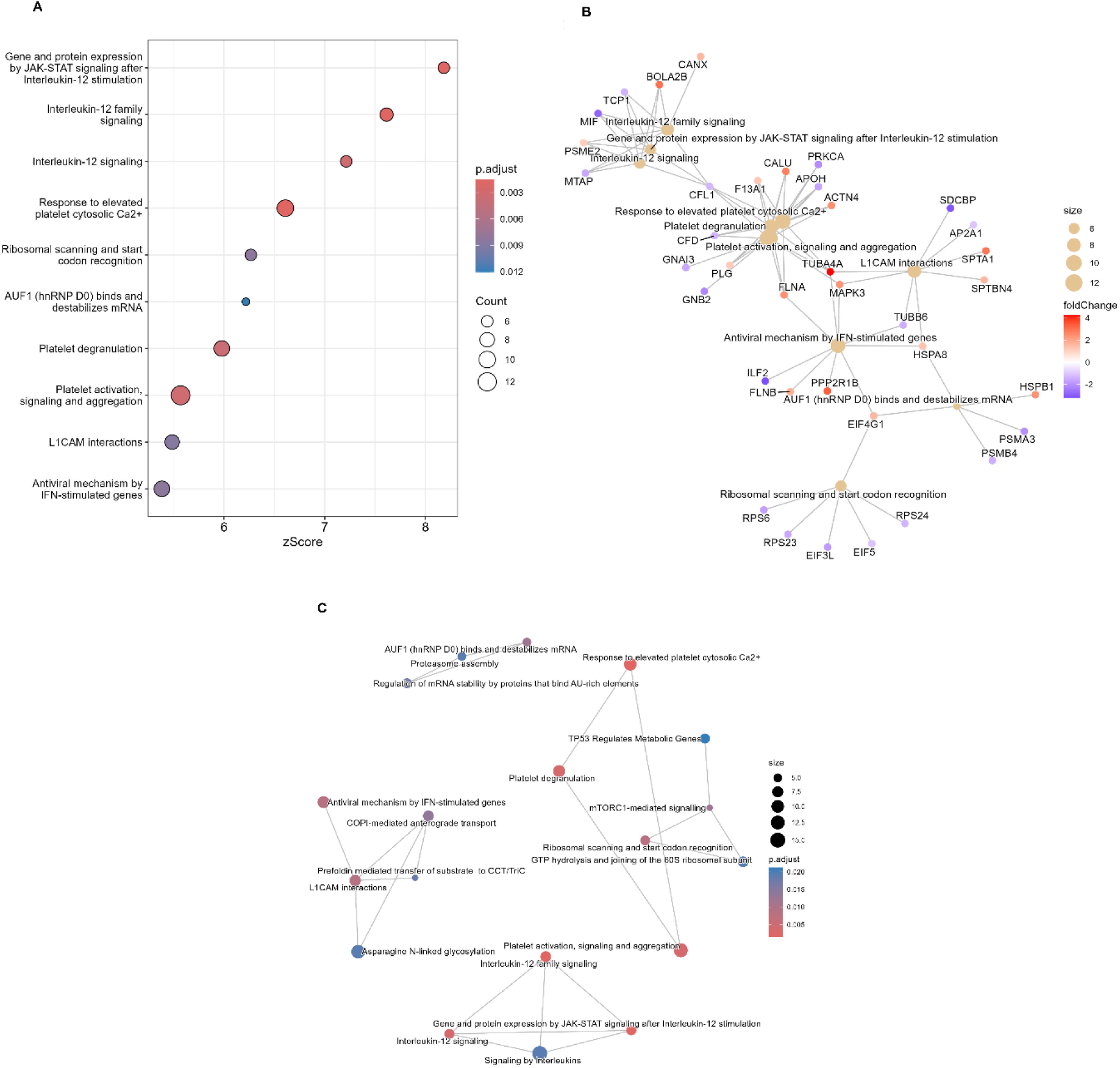
Reactome pathway enrichment analysis highlights immune signaling and translational control alterations in AF samples. (A) Dot-plot showing significantly enriched Reactome pathways from over-representation analysis of differentially abundant proteins from AF-induced left atrial tissue versus sham-operated controls. The x-axis represents enrichment z-score, dot size indicates gene count, and colour intensity represents adjusted p-value (Benjamini-Hochberg correction). Gene and protein expression by JAK-STAT signalling and interleukin-12 family signalling pathways show the most significant enrichment. (B) Concept Network Enrichment (CNE) plot displaying gene-concept relationships for top enriched Reactome pathways. Genes (outer nodes) are connected to their associated pathway terms (inner nodes), with node size proportional to gene count or term size. Gene nodes are coloured by log2-fold change (red = upregulated, blue = downregulated in AF). (C) Enrichment Map Analysis (EMA) plot showing functional relationships between enriched Reactome pathways. Nodes represent pathway terms sized by gene count, and edges connect pathways sharing genes. Terms cluster into distinct functional modules including ribosomal scanning/codon recognition, platelet activation/degranulation, and interleukin-12 signalling networks. All enrichments shown have adjusted p-value < 0.05.

### Consensus analysis

GO, KEGG and WikiPathways use a broadly similar database structure and so next we searched for consensus of enrichment amongst these three databases. Only one pathway was conserved between all three databases, Oxidative Phosphorylation with median *p-value* of 8.7 e-4. The consensus analysis is represented graphically in Figure 7 with an UpSet graph.

**Figure 7.**
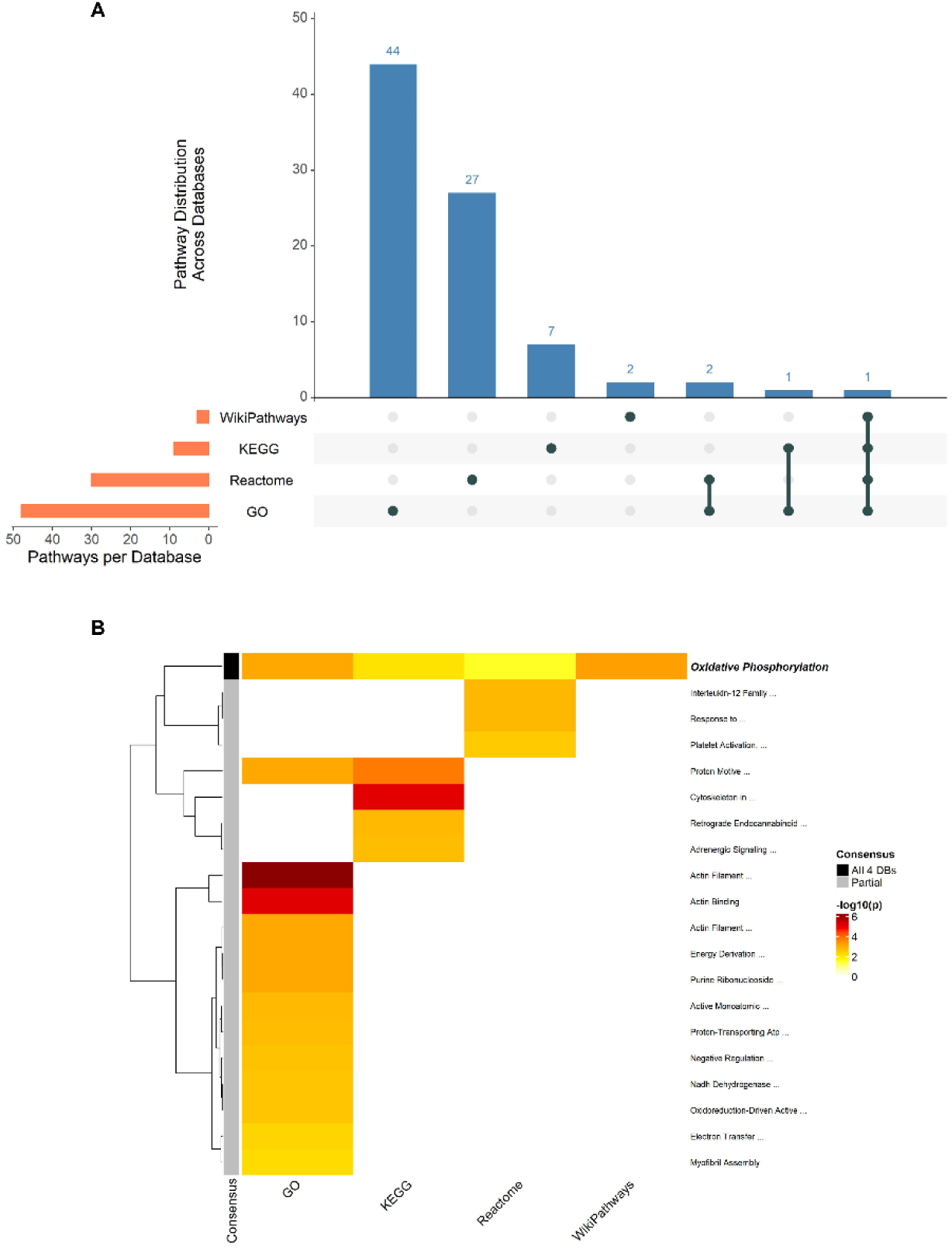
Multi-database pathway enrichment analysis reveals consensus on the Oxidative Phosphorylation pathway. (A) UpSet plot displaying the distribution of enriched pathways across Gene Ontology (GO), KEGG, and WikiPathways databases. Vertical bars indicate the number of pathways found in each database combination (intersections), while horizontal bars show the total number of enriched pathways per database. Pathways were considered significantly enriched at adjusted p-value < 0.01. note only one pathway is enriched in all three databases. (B) Heatmap of pathway enrichment significance across databases. Colour intensity represents -log10(adjusted p-value), with darker red indicating higher statistical significance. Each row represents a unique pathway description (lowercase-matched across databases). The top consensus pathway, appearing in multiple databases with the strongest median significance, is labelled (Oxidative Phosphorylation). Columns represent individual enrichment databases; rows are unsupervised hierarchically clustered.

### Mitochondrial specific analysis

Having established that unbiased enrichment data analyses broadly support the enrichment of mitochondrial proteins, we moved to mitochondrial specific analysis using the MitoCarta database and Mitology R-package. Gene set enrichment analysis (GSEA) on the MitoCarta database and using all data with log2fold change weighted *p-value* as the metric revealed only one significantly enriched cluster, OXPHOS subunits, i.e. proteins involved with OXPHOS. We show the GSEA plot, CNE and ranked-gene scatter plot in Figure 8 (A), (B) and (C) respectively. OXPHOS subunits are dysregulated with Normalized Enrichment Score (NES)/ zScore 1.76 and BH adjusted *p-value* of 0.016.

**Figure 8.**
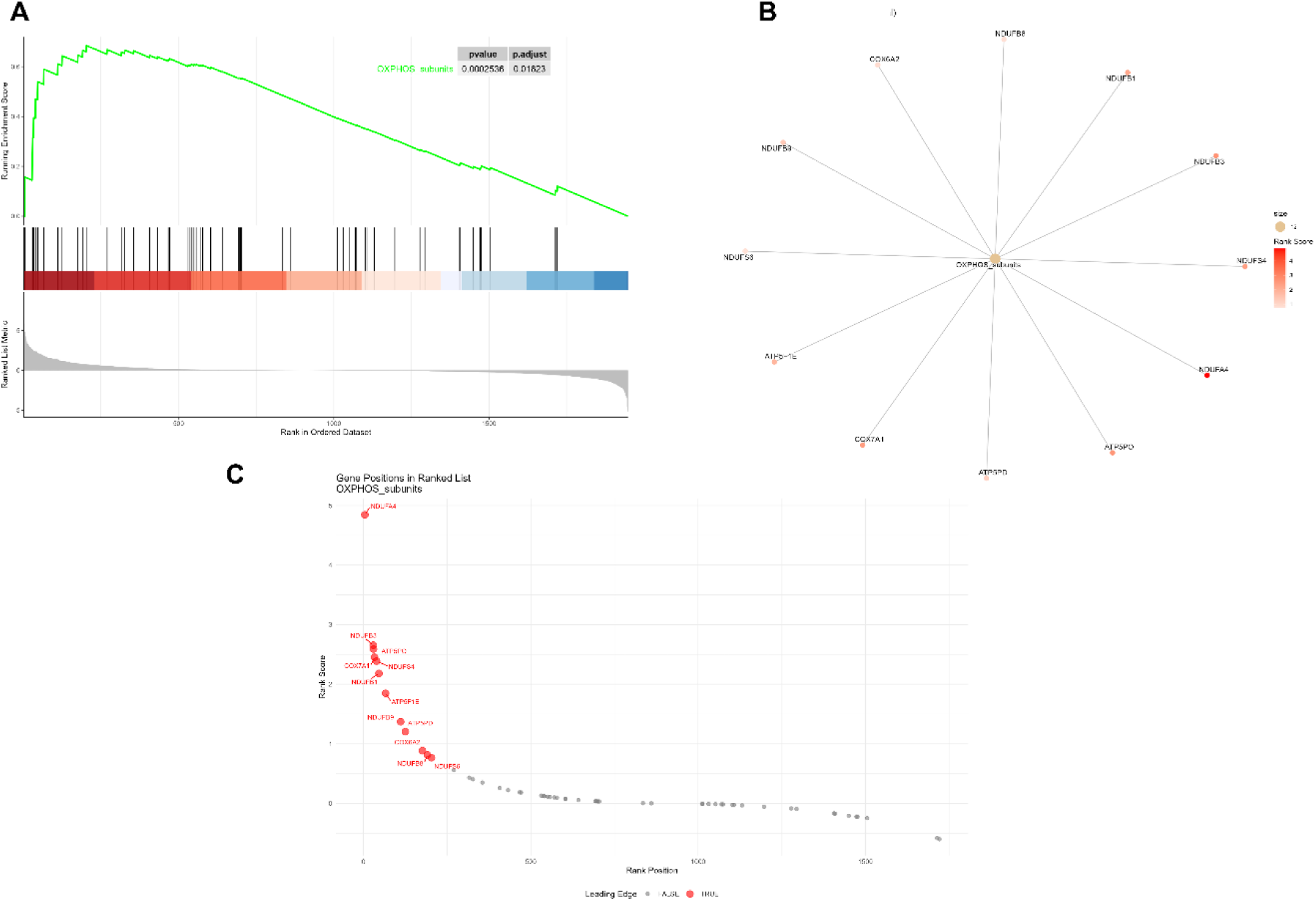
Mitochondrial-specific protein set enrichment analysis reveals upregulation of oxidative phosphorylation subunits in AF samples. (A) Gene Set Enrichment Analysis (GSEA) plot for the OXPHOS subunits pathway from the MitoCarta database. The top panel shows the running enrichment score (green line) across the ranked protein list, with vertical black lines indicating the position of genes within the OXPHOS subunits protein set. The bottom panel displays the ranked list metric (log2-fold change weighted p-value), with red-to-blue colouring indicating upregulated to downregulated genes in AF versus sham-operated samples. The enrichment score peaks early in the ranked list, indicating significant upregulation of OXPHOS subunits (NES = 1.76, p-value = 0.00025, adjusted p-value = 0.016). (B) Concept Network Enrichment (CNE) plot showing gene-concept relationships for the OXPHOS subunits pathway. Individual proteins (outer nodes) are connected to the central OXPHOS subunits pathway node, with node size proportional to gene count (n=12). Protein nodes are coloured by rank score, with darker red indicating higher-ranked (more significantly upregulated) genes in AF samples. Key NADH dehydrogenase (NDUF) subunits, ATP synthase components (ATP5), and cytochrome c oxidase (COX7A1) are displayed. (C) Gene position plot showing the ranked positions of OXPHOS subunits proteins within the ordered dataset. The x-axis represents rank position in the full gene list, and the y-axis shows the rank score. Leading edge proteins (contributing to the enrichment signal) are highlighted in red, while non-leading-edge proteins are shown in grey. OXPHOS subunit genes cluster prominently at high ranks (left side), with NDUFA4 showing the highest rank score.

### Network analysis reveals HSPA9-mediated coordination of OXPHOS complex assembly in AF-induced atrial mitochondria

To move beyond pathway enrichment and identify potential causal mechanisms underlying mitochondrial dysfunction in AF, we leveraged the directional regulatory relationships encoded in the Reactome pathway database. GSEA using the MitoLogy package confirmed OXPHOS subunits as the most significantly enriched pathway in the mitochondrial proteome of AF-induced left atrial tissue compared to sham-operated controls (NES = 1.8, FDR = 0.04; Figure 9). We then extracted the “Aerobic respiration and respiratory electron transport” pathway from Reactome as a directed graph, which contained 255 mitochondrial proteins connected by 6,779 edges encoding physical interactions and regulatory relationships.

**Figure 9.**
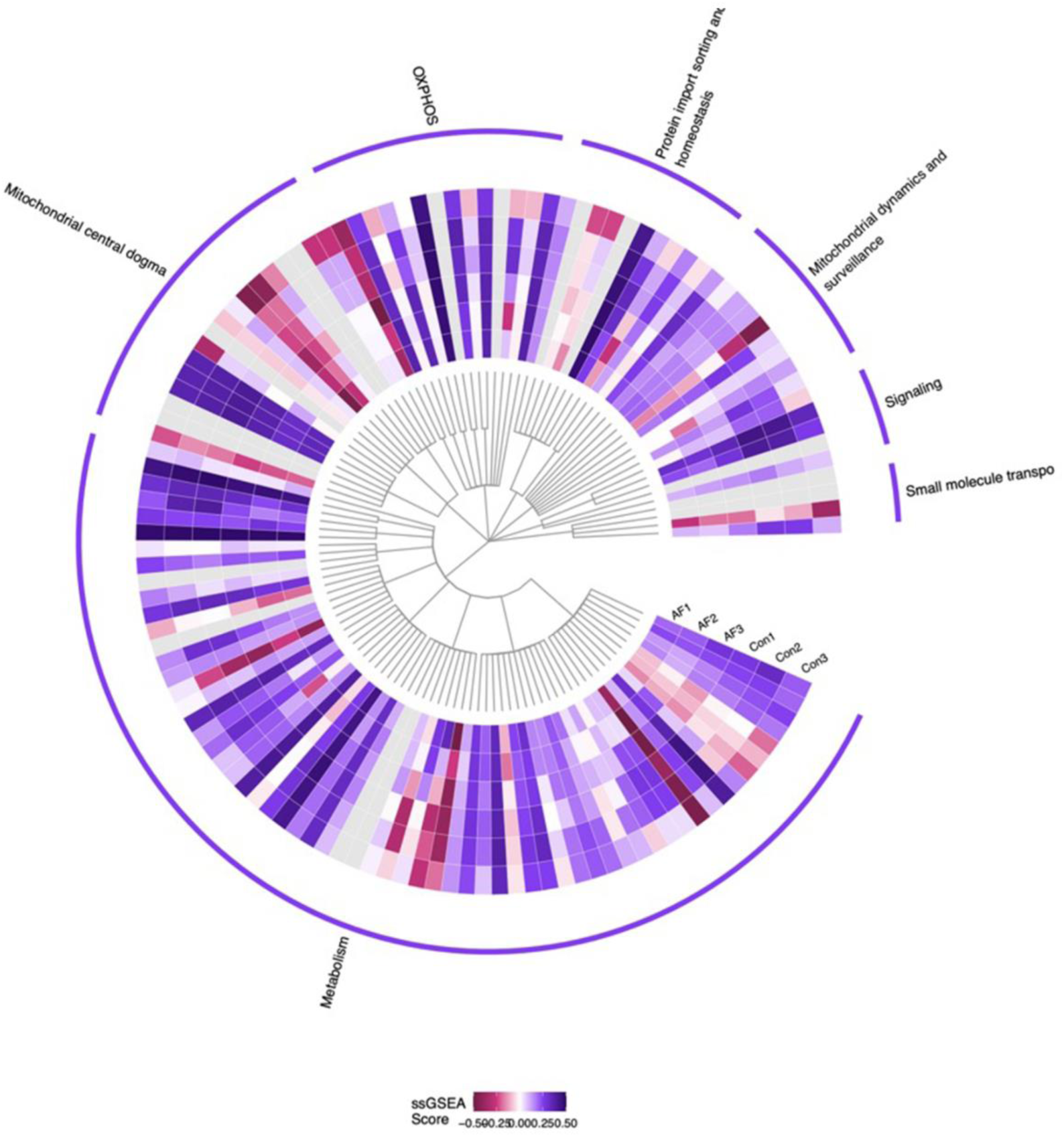
Comprehensive mitochondrial pathway analysis reveals functional remodelling across major mitochondrial processes in AF samples. Heatmap displaying single-sample Gene Set Enrichment Analysis (ssGSEA) scores for hierarchically organized mitochondrial pathways from the MitoCarta database. The central phylogenetic tree represents the hierarchical relationships between mitochondrial gene sets, organized into seven major functional categories (indicated by purple arc labels): OXPHOS, Protein import sorting and homeostasis, Mitochondrial dynamics and surveillance, Signalling, Small molecule transport, Mitochondrial central dogma, and Metabolism. Each concentric ring represents an individual sample (AF1, AF2, AF3 = AF-induced left atrial tissue; Con1, Con2, Con3 = sham-operated controls). Colour intensity represents ssGSEA enrichment scores, with red indicating pathway upregulation, purple indicating pathway downregulation, and white indicating no enrichment (score range: -0.50 to +0.50). Each radial segment corresponds to a specific mitochondrial pathway or gene set, with related pathways clustering together based on functional similarity. The visualization reveals coordinated patterns of mitochondrial pathway activity across samples, with notable enrichment of OXPHOS-related pathways in AF samples compared to controls.

By overlaying our differential expression data onto this regulatory network (Table 1), we identified 94 regulatory edges (activation or inhibition relationships) where both the regulator and target proteins were quantified in our dataset. Strikingly, 69.1% (*n*=65) of these regulatory relationships showed concordance, that is, proteins in activation relationships changed in the same direction, while proteins in inhibition relationships changed in opposite directions (median concordance score: 0.99). This concordance rate significantly exceeds that arising from random permutation of the data (59.9%, 95% CI: 52.1%-67.0%, 1000 permutations, *p-value* = 0.017).

**Table 1.** Composition of the OXPHOS causal regulatory network in AF samples: Summary of proteins involved in the regulatory network derived from the Reactome “Aerobic respiration and respiratory electron transport” pathway. Proteins are categorized by their functional classification within the oxidative phosphorylation system. The network shows preferential enrichment of Complex I subunits, consistent with HSPA9-mediated coordination of NADH dehydrogenase assembly. Pathway Concordance represents the proportion of regulatory edges (activation or inhibition relationships) showing directionally consistent changes between regulator and target proteins, significantly exceeding that expected by chance (permute test). Concordance: 65/94 regulatory edges (69.1% vs. 59.8% expected, *p* = 0.017).

| Component | Number of Proteins |
| --- | --- |
| Complex I subunits | 26 |
| Complex III subunits | 5 |
| Complex IV subunits | 1 |
| Chaperones/Import machinery | 2 |
| Other OXPHOS-related | 25 |
| Total unique proteins | 59 |

To identify key regulatory nodes driving this coordinated response, we searched for multi-step causal chains where protein A activates B, and B activates C, with all changes directionally consistent. This analysis revealed 22 three-step cascades. Network topology analysis identified HSPA9 (mitochondrial heat shock protein 70) as the primary regulatory hub, serving as an intermediate node in 7 of these chains, followed by TIMM21 (*n*=4), UQCRQ (*n*=3), and UQCRFS1 (*n*=2).

The constructed network centred on HSPA9 revealed a striking regulatory architecture (Figure 10): HSPA9 and Rieske iron-sulphur protein, Complex III (UQCRFS1) (Rieske iron-sulphur protein, Complex III) have reciprocal activation, (chain concordance score: 10.5). This suggests that HSPA9 then coordinates the dysregulation of multiple Complex I subunits including NDUFV2, NDUFV1, NDUFS1, NDUFS2, NDUFS7, and NDUFS8 (chain concordance scores: 5.7-10.5).

**Figure 10.**
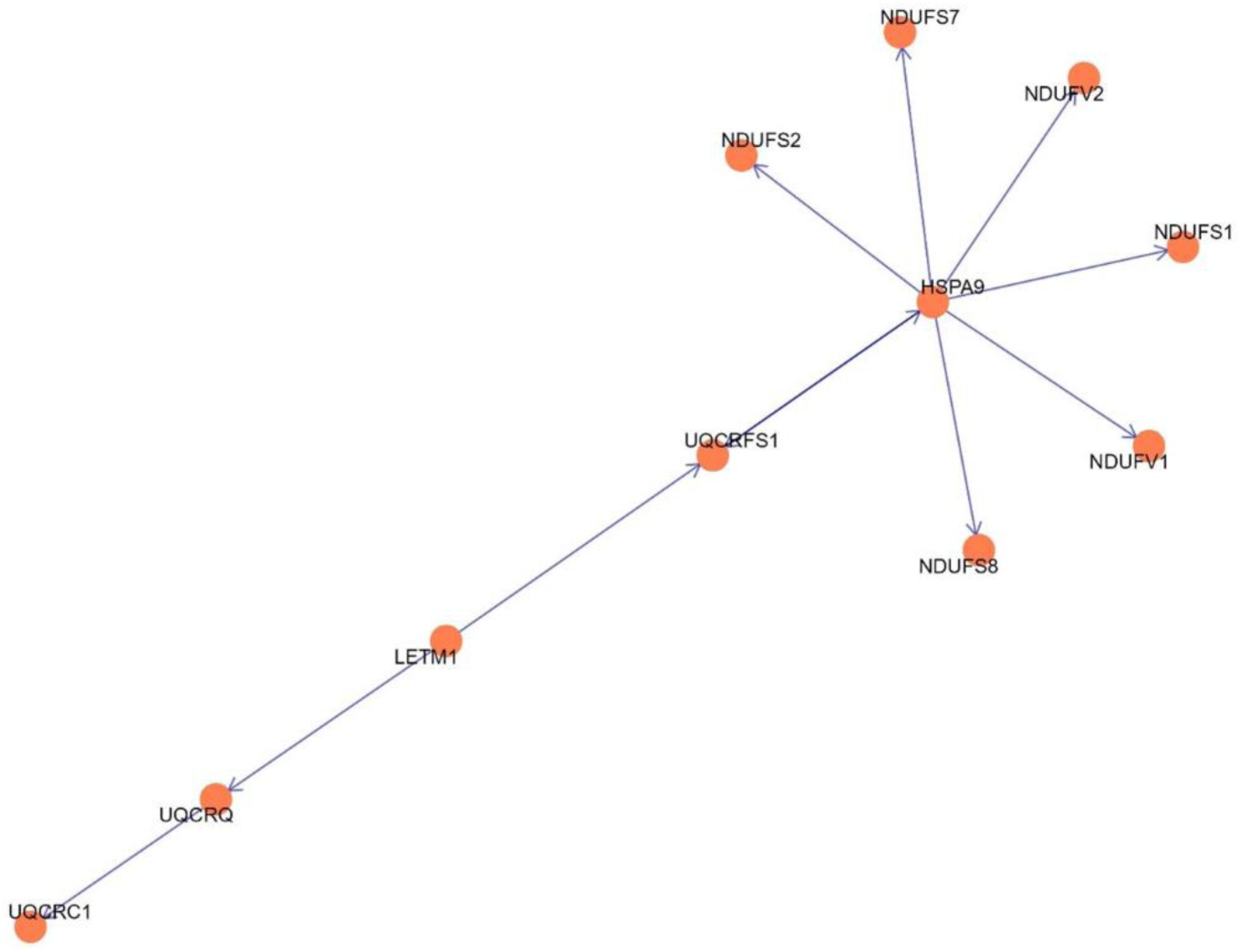
Inferred regulatory network reveals HSPA9 as a master coordinator of OXPHOS complex assembly in AF samples. Directed network graph displaying multi-step causal cascades within the OXPHOS pathway showing high concordance between regulator and target protein expression changes. The network was constructed by overlaying differential abundance data onto directional regulatory relationships from the Reactome “Aerobic respiration and respiratory electron transport” pathway. Nodes represent proteins identified in AF-induced left atrial tissue versus sham-operated controls, with all proteins shown in orange indicating upregulation in AF samples. Node size reflects the protein’s hub centrality in the network architecture. Directed edges (arrows) represent activation relationships derived from Reactome regulatory annotations. HSPA9 (mitochondrial eat shock protein 70, mt-Hsp70) emerges as the primary regulatory hub, participating in 7 of 22 identified three-step directional cascades. HSPA9 forms a reciprocal activation loop with UQCRFS1 (Rieske iron-sulphur protein, Complex III) and coordinates the expression of multiple NADH dehydrogenase subunits (NDUFV1, NDUFV2, NDUFS1, NDUFS2, NDUFS7, NDUFS8). The cascade extends through UQCRFS1 to downstream Complex III components (LETM1, UQCRQ, UQCRC1), revealing a hierarchical regulatory architecture. All displayed relationships show directional concordance (activation relationships with co-directional changes), with chain concordance scores ranging from 5.7 to 10.5.

## Discussion

This study aimed to uncover mechanisms underlying mitochondrial dysfunction (mitochondrial fractions purified from goat) in AF using a bioinformatic approach. Analysis of the mitochondrial proteome from AF-induced left atrial tissue revealed HSPA9 as a central regulatory hub coordinating OXPHOS complex assembly, with preferential enrichment of Complex I subunits. Network mapping identified 94 regulator–target relationships, with statistically significant pathway concordance, activation pairs changed in the same direction, indicating system-wide, coordinated regulation of the electron transport chain rather than independent responses to AF. Notably, the network centered on HSPA9 as a regulatory hub for OXPHOS proteins, reciprocally interacting with UQCRFS1 (Complex III), suggesting a striking regulatory architecture driving mitochondrial remodelling in AF.

The broader network encompassed 26 Complex I proteins, 5 Complex III proteins, 1 Complex IV protein, and 2 protein import/chaperone factors (Table 1), suggesting HSPA9-mediated coordination of respiratory chain complex assembly as a central mechanism in AF-induced mitochondrial remodelling. Notably, the 69.1% concordance across the OXPHOS regulatory network (65 of 94 regulatory edges) significantly exceeded the 59.8% expected from permutation testing that preserved the observed fold-change distribution (95% CI: 52.1–67.0%, *p*-*value* = 0.017). This finding suggests that the observed changes are not random but represent system-wide, coordinated regulation of the electron transport chain in response to AF. The predominance of Complex I subunits within this network suggests targeted remodelling to optimize electron flux and maintain mitochondrial function under AF-induced stress, reinforcing the role of HSPA9 as a key regulatory hub driving these adaptations.

AF is a multifactorial arrhythmia characterized by electrical instability, structural remodelling, and metabolic stress [37–39]. Recent proteomic studies, including compartmentalization analyses, have revealed that AF involves profound remodelling of mitochondrial and EL networks. In particular, Ayagama *et al.* (2024) demonstrated that AF in goats disrupts EL protein trafficking and calcium signalling, implicating organelle crosstalk in disease progression [26]. Building on this, analysis of our mitochondrial fraction proteomics study provides a systems-level view of AF-associated molecular disruptions, highlighting consistent patterns of mitochondrial dysfunction, cytoskeletal reorganization, and immune-metabolic signalling. Electrical remodelling and the resulting arrhythmogenic substrate in AF may be driven, in part, by cellular energy deficits and oxidative stress stemming from mitochondrial dysfunction [40]. While the precise role of mitochondrial impairment in AF pathogenesis remains incompletely understood, its importance is underscored by observations that therapies enhancing mitochondrial function can reduce the burden of AF [41]. Together, these findings suggest that AF is accompanied by organelle stress responses, altered energy metabolism and structural instability within atrial tissue.

### Mitochondrial dysfunction and energetic remodelling

Across multiple analyses, OXPHOS emerged as a robust and recurrently enriched pathway. Figure 4 indicates mitochondrial dysfunction and energetic remodelling, with coordinated changes in OXPHOS driven largely by altered Complex I related pathways and genes. This likely reflects elevated energy demands in AF-affected atrial tissue, where impaired contractility and electrical signalling require metabolic adaptation [42, 43].

Previous studies have shown that mitochondrial dysfunction contributes to AF by impairing ATP synthesis and increasing ROS, which destabilize ion channels and gap junctions [42, 43]. The enrichment of ATPase activity and mitochondrial protein folding pathways, particularly involving *HSPA9*, points to stress-induced remodelling of the respiratory chain. Directed network analysis revealed *HSPA9* as a master regulator of Complex I and III subunits, reinforcing its role in maintaining mitochondrial integrity under pathological conditions. HSPA9 downregulation has been shown to diminish peroxisome abundance and augment autophagy by promoting SQSTM1-dependent Pexophagy, a mechanism relevant to cardiac pathology given that peroxisomal dysfunction and disrupted autophagic flux are key drivers of ischemic injury, cardiomyopathy and heart failure [44]. Our AF study shows HSPA9 functions as a mitochondrial chaperone essential for maintaining cardiomyocyte viability under oxidative stress suggesting the importance of *HSPA9* in cellular regulation and disease.

### Cytoskeletal Reorganization and Structural Instability

Proteomic changes showed alterations in key cytoskeletal processes, filament bundle formation and extracellular matrix organisation. These shifts indicate that atrial myocytes are undergoing structural remodelling, which can interfere with how they sense and respond to mechanical forces and may promote fibrosis. Disruption of the cytoskeleton has also been associated with uneven electrical conduction and the development of arrhythmogenic substrates in AF [45, 46].

### Immune activation and stress signalling

Figure 6 shows evidence of immune-related and stress-response signalling. Enriched pathways in Panel A include complement activation and cytokine-associated processes, while Panels B and C highlight networks involving complement proteins and heat-shock–associated stress genes. Together, these patterns indicate that innate immune pathways and cellular stress responses are activated, reflecting a state where atrial tissue is responding to inflammation, protein misfolding, and environmental stress signals. These findings suggest that AF is accompanied by inflammatory signalling and proteotoxic stress, which may exacerbate tissue remodelling and electrical instability [47].

### Translational and post-transcriptional regulation

Enrichment of ribosomal scanning, TP53-regulated metabolic genes, and spliceosomal pathways points to deeper layers of molecular reprogramming. These changes may reflect altered protein synthesis and RNA processing in response to chronic stress [48, 49], further contributing to the maladaptive phenotype observed in AF. Notably, ER stress–associated autophagy has been identified as a critical pathway in AF progression, suggesting that the observed translational and spliceosomal reprogramming maybe linked to disrupted ER stress responses and impaired proteostasis [49].

### Integrated model of AF pathogenesis

Taken together, the data support a model in which AF causes mitochondrial stress, cytoskeletal disruption, and immune-metabolic imbalance (Table 2). The proteomic landscape reveals both adaptive and maladaptive responses, with mitochondrial compensation attempting to sustain energy demands while structural and inflammatory changes destabilize atrial function.

**Table 2:** Summary of key pathophysiological axes identified in goat atrial fibrillation (AF) based on mitochondrial fraction proteomics. Each axis represents a distinct biological domain disrupted in AF, with associated molecular features derived from enrichment analyses, network modelling, and gene set profiling. These findings highlight the convergence of mitochondrial dysfunction, structural remodelling, immune activation, and translational reprogramming in the disease process.

| Pathophysiological Axis | Key Features |
| --- | --- |
| **Mitochondrial Dysfunction** | Impaired oxidative phosphorylation, chaperone failure, energy imbalance |
| **Cytoskeletal Remodelling** | Actin filament disruption, ECM changes, altered mechano-transduction |
| **Immune Activation** | Complement system, JAK-STAT signalling, heat shock protein response |
| **Translational Reprogramming** | TP53-mediated metabolic regulation, ribosomal and spliceosomal shifts |

These insights not only align with known mechanisms of AF but also reveal novel mitochondrial regulators and network vulnerabilities that may serve as therapeutic targets. Future studies should determine whether this remodelling enhances resilience or predisposes to mitochondrial instability under chronic AF and assess the translational relevance of these findings in human AF, including testing whether modulating mitochondrial function or cytoskeletal integrity can restore atrial stability.

### Study limitations

The mitochondrial bioinformatics analyses in this study are based on protein abundance data integrated with published databases and pathway annotations. While these analyses provide insights into mitochondrial remodelling and energetic adaptation in AF, they represent inferred relationships rather than experimentally validated causal mechanisms. Establishing definitive pathway alterations will require additional functional and mechanistic studies.

Proteomic measurements were obtained from atrial tissue collected at a single time point in a chronic AF goat model. Consequently, the results provide a snapshot of mitochondrial protein regulation, which may not capture dynamic changes occurring during disease onset, progression from paroxysmal to persistent AF, or during acute AF episodes.

While the goat AF model recapitulates many features of human AF, including structural, electrical, and metabolic remodelling, it may not fully replicate the heterogeneity and multifactorial nature of the human disease. Differences in species-specific mitochondrial regulation, cardiac metabolism, or progression of AF may limit the direct translation of findings to humans.

The mitochondrial fraction analysed in this study, while enriched, may not capture all mitochondrial proteins or transient protein complexes, particularly low-abundance or highly dynamic components. Consequently, some relevant mitochondrial pathways or regulatory mechanisms may remain underrepresented.

While this study focused on proteomics, mitochondrial function is also influenced by transcriptomic, metabolomic, and post-translational modifications. The lack of concurrent multi-omics measurements limits the ability to fully integrate regulatory mechanisms or temporal responses of mitochondrial pathways in AF.

## Conclusions

Our previous work [26] investigated the proteomic landscape of left atrial tissue in a goat model of AF, analysing both TL and EL fractions. That study highlighted dysregulation of multiple pathways relevant to mitochondrial function, including OXPHOS, AMPK signalling, autophagy, protein folding, and NADPH oxidase-mediated ROS generation. Notably, in that study [26], AMPK was observed to be upregulated in response to ATP depletion, consistent with elevated energetic demands in AF atrial tissue. Increased ATP consumption was linked to maintenance of cellular ion homeostasis, while dysregulated lysosomal V-ATPase activity further indicated organelle-specific contributions to energy stress. Dysregulation of protein folding and chaperone pathways suggested an active proteostatic response to maintain mitochondrial and cellular function under stress conditions.

Building on these observations, our study focused specifically on the mitochondrial fraction, using detailed network analyses to examine energetic remodelling in AF. Consistent with recent findings in goat AF [26], we observed upregulation of key mitochondrial subunits. Across the results, mitochondrial representation is dominated by Complex I subunits, including NDUFS1, NDUFS2, NDUFS4, NDUFS7, NDUFS8, NDUFV2, NDUFA2, NDUFB3 and NDUFB6, which appear prominently in Figures 4B and 10. Complex III components (UQCRC1, UQCRQ, UQCRFS1) and the mitochondrial chaperone HSPA9, together with LETMD1/LETM1, are also evident (Figure 10), while ATP5PF is the only detectable Complex V subunit (Figure 4B). Collectively, the visual evidence, particularly from Figures 4B and 10, indicates an OXPHOS profile heavily skewed toward Complex I involvement, with additional contributions from Complex III and limited Complex V representation.

Our data also suggested HSPA9 could be a central regulator of Complex I and III subunits, supporting a role for chaperone-mediated maintenance of respiratory chain integrity under pathological conditions. Furthermore, our analyses reinforced the connection between mitochondrial dysfunction, ROS generation, and downstream effects on ion channel and gap junction stability.

Pronto *et al.* [50] demonstrate that impaired mitochondrial Ca^2+^ handling and resultant energetic dysregulation contribute to ROS generation and arrhythmogenic triggers in AF. Complementing these findings, our proteomic analyses of the mitochondrial fraction in the goat AF model reveal dysregulation of OXPHOS subunits, and chaperone-mediated maintenance of the respiratory chain. Together, these studies show that mitochondrial adaptation and dysfunction are prominent features of the energetic and electrophysiological remodelling observed in AF. Although rapid atrial activation, increased workload and elevated energy demands are well recognized in AF, the contribution of mitochondria, the main energy producing organelles in cardiac myocytes, to the onset and persistence of AF remains poorly defined [40, 51]. Recent studies [26], together with our current findings, indicate that AF in the goat model is accompanied by mitochondrial stress, energetic remodelling and adaptive proteostatic responses. These changes occur alongside broader alterations in cytoskeletal organisation and immune-metabolic signalling, suggesting that mitochondrial adjustments reflect the heightened energetic burden of AF, while structural and oxidative stress are part of the atrial remodelling associated with the arrhythmogenic substrate. Importantly, these mitochondrial proteomic data add a novel organelle-specific layer to the existing goat AF dataset, enhancing mechanistic insight and providing a valuable resource for future studies on AF pathophysiology.

## Acknowledgements

This study was supported by the Wellcome Trust and Royal Society (PI RABB: 109371/Z/15/Z). RABB is funded by the Wellcome Trust and British Heart Foundation (109371/Z/15/Z). RABB acknowledges research funds from the Ellis T Davies Fellowship Endowment, University of Liverpool. RABB held a Senior Research Fellowship at Linacre College). This research was funded in whole, or in part, by the Wellcome Trust [109371/Z/15/Z]. For the purpose of Open Access, the author has applied a CC BY public copyright license to any Author Accepted Manuscript version arising from this submission. We thank Dr Rebecca A Capel for support in tissue preparation. This work was also supported by the Netherlands Heart Foundation (EmbRACE: Electro-Molecular Basis and the theRapeutic management of Atrial Cardiomyopathy, fibrillation and associated outcomEs), the European Union (MAESTRIA: Machine Learning Artificial Intelligence Early Detection Stroke Atrial Fibrillation, grant number 965286) and by the Leducq Foundation (Immune Targets for the treatment of atrial fibrillation).

## Authorship contribution statement

RABB: Conceptualization and reviewing; RABB, RBJ: Investigation, writing the first draft, reviewing and editing. RABB: Fund acquisition via the Wellcome Trust Fellowship; US and SV: Fund acquisition via Netherlands Heart Foundation, European Union, Leducq Foundation. TA, RABB and RBJ: Lead experimentalists, formal analysis, writing, reviewing and editing – original draft. RABB, US, SV: Supervision, data discussion. RF, SH: Conducting MS/MS studies. US, SV: Goat AF model, physiology experiments and tissue collection.

## Data Availability

Original Mitochondrial data were submitted to PRIDE (PRIDE ID: PXD041056). Please contact the corresponding authors for information and data related to the manuscript.

## Supplementary Files

**Supplementary File 1:** Comprehensive list of quantified mitochondrial fraction proteins identified in the data for *Capra hircus* (goat) including -log(*p-values*) and LFC.

